# Moxifloxacin efficacy against *Mycobacterium abscessus in vivo* using zebrafish model

**DOI:** 10.1101/865824

**Authors:** Wenjuan Nie, Shan Gao, Tianlu Teng, Wenqiang Zhou, Yuanyuan Shang, Wei Jing, Wenhui Shi, Qingfeng Wang, Xuerui Huang, Baoyun Cai, Jun Wang, Jing Wang, Ru Guo, Qiping Ge, Lihui Nie, Xiqin Han, Yadong Du, Naihui Chu

## Abstract

Moxifloxacin (MFX) showed good activity *in vitro* against *Mycobacterium abscessus* (*M. abscessus*) and was suggested as one of the antibiotic regimens for adults with *M. abscessus* disease. However, some other studies showed that MFX showed less or none activity against *M. abscessus*. In our study we aim to evaluate MFX activity against *M. abscessus* using zebrafish (ZF) model *in vivo*. MIC of each drugs were determined by broth microdilution method. *M. abscessus* labeled by CM-DiI, were micro-injected into ZF. Survival curves were determined by recording dead ZF every day. After 4 days of incubation ZF were lysed. Colony-forming unit (CFU) were enumerated and results are expressed as mean log10 CFU per ZF. Bacteria dissemination and fluorescence intensity in ZF were observed and analyzed. Inhibition rate was also calculated. In our study MFX showed good activity *in vitro*. But *in vivo* MFX showed limited restriction to *M. abscessus*. The association between increased survival and high dose of MFX is not significant. Same results were observed in bacterial fluorescence intensity and inhibition rates, with no significant difference when compared with no drug group (P > 0.05). However, significant difference was observed in azithromycin (AZM) group. MFX showed limited efficacy on *Mycobacterium abscessus in vivo* using ZF model. MFX’s activity *in vivo* need to be confirmed.

## Introduction

Nontuberculous mycobacterial (NTM) disease in humans has been recognized as an emerging public health problem [1,2]. Drug therapy of NTM disease is long, costly, and often associated with drug-related toxicities, so the treatment of NTM disease can be disappointing. Clinical improvement and prolonged culture conversion are not achievable for all patients [3]. There are two kinds of NTM: rapidly growing mycobacteria (RGM) and slowly growing mycobacteria (SGM). *Mycobacterium abscessus* (*M. abscessus*) is the most common etiological agent of disease caused by rapidly growing mycobacteria [4–7]. The emerging pathogen *M. abscessus* is the etiological agent of a wide spectrum of infections in humans, including severe chronic pulmonary and disseminated infections [8]. With current antibiotic options, *M. abscessus* is a chronic incurable infection for most patients. At present, there is no reliable or dependable antibiotic regimen, even based on *in vitro* drug susceptibility testing (DST) and including parenteral agents, to produce cure for *M. abscessus* disease [4]. *M. abscessus* infections may lead to outbreaks of infection [9, 10]. Previously, *M. abscessus* were thought to be independently acquired by susceptible individuals from the environment. However, using whole-genome analysis of a global collection of clinical isolates, it shows that the majority of *M. abscessus* infections are acquired through transmission, potentially via fomites and aerosols, of recently emerged dominant circulating clones that have spread globally. This represents an urgent international infection challenge [11].

The major issue with *M. abscessus* relies on its intrinsic resistance to the most available antibiotics. The American Thoracic Society has recommended different groups of agents, namely, macrolides (clarithromycin), aminoglycosides (amikacin), cephamycins (cefoxitin), and carbapenems (imipenem), to treat *M. abscessus* infections [4]. Moxifloxacin (MFX) emerged as pnromising candidates for the treatment of RGM infections [4,12,13]. It showed good activity *in vitro* against *M. abscessus* [14] and was suggested as one of the antibiotic regimens for adults with *M. abscessus* disease [12]. However, some other studies showed that MFX showed less or none activity against *M. abscessus in vitro* [15-18]. DST *in vitro* may be necessary, but DST is not fully standardized. Further more, the clinical response to drugs does not correlate well with *in vitro* DST. Future work should also address MFX efficacy *in vivo* [19]. This further emphasizes the need for suitable animal models [20,21]. Recently, the *M. abscessus*/zebrafish (ZF) model already provided important insights into infectious diseases pathogenesis. It is rapidly gaining favor as a useful model for the study of host–bacterial interactions [22-25]. Because of its genetic tractability and optical transparency, ZF represent an exquisite model to study many aspects of *M. abscessus*. Such a simple and innovative system would be particularly suited to assess antibacterial activities for the discovery of the urgently needed drugs to fight *M. abscessus* [19].

In this study, we reported experimental conditions *in vivo* imaging of *M. abscessus* infections and their use to test the efficacy of drug treatments. We aim to evaluate MFX activity against *M. abscessus* using ZF model, a system that could be applied to high-throughput *in vivo* testing of drug efficacy against the most drug-resistant mycobacterial species, so as to make certain MFX’s confusing role for *M. abscessus* treatment.

## Materials and Methods

### 1. Minimum inhibitory concentration(MIC)

The standard isolate ATCC19977 was used for culture. Isolate was subcultured on the Lowenstein-Jensen medium at 37°C for about 4-6 days to observe colony morphology. MFX and azithromycin (AZM) were purchased from Sigma–Aldrich Chemical Co. Drugs were solved according to the Clinical and Laboratory Standards Institute (CLSI). The final drug concentrations of MFX and AZM were 0.0625 to 32 μg/ml and 0.5 to 256 μg/ml, respectively. MIC of each drugs were determined by broth microdilution method as CLSI recommended, using 96-well plates. The MIC was determined on after 3 days’ incubation. After incubation, Alamar blue dye (Serotec) was added to each well and the plates were reincubated for 24h. A color change from blue to pink indicated bacterial growth. MIC was defined as the lowest concentration of the drug that showed no color change, which were the lowest concentrations of drugs capable of inhibiting the visible growth of tested isolates. DST results were evaluated according to CLSI breakpoints recommendations.

### 2. Microinjection of *M. abscessus* into ZF

ZF experiment was approved by Ethics Committee in Beijing Chest Hospital affilicated to Capital Medical University.

We adapted a previously designed protocol [19] to assess the activity of MFX and AZM against *M. abscessus* in ZF. *M. abscessus* ATCC19977 with a smooth morphotype were grown in Middlebrook 7H9 broth (Becton Dickinson) supplemented with 10% OADC (Becton Dickinson) and 0.05% Tween 80 (Sigma-Aldrich) at 30°C. Mid-log-phase cultures of *M. abscessus* were centrifuged, washed, and resuspended in PBS supplemented with 0.05% Tween 80. Bacterial suspensions were then homogenized and sonicated, and the remaining clumps were allowed to settle for 5 to 10 min, as previously described [26]. Bacteria were concentrated to an optical density in PBS and i.v. injected. *M. abscessus* labeled by red fluorescent CM-DiI, were micro-injected into 3 days post-fertilization (dpf) wide type ZF caudal vein. Different concentration and amounts of bacteria, and different observation duration were tested for establishing infected ZF model.

### 3. Maximum tolerance concentrations (MTC) of MFX and AZM in ZF model *in vivo*

10 3dpf ZF without *M. abscessus* infection were randomly selected and placed into 1 well of 24-well plates, with each well of having 1ml water. MFX or AZM was then directly added to the water. MFX with concentrations of 10μg/ml, 100μg/ml, 250μg/ml, 500μg/ml, 1000μg/ml, 2000μg/ml, and AZM concentrations of 1μg/ml, 10μg/ml, 100μg/ml, 250μg/ml, 500μg/ml and 1000μg/ml were tested, separately. Drug-containing water was renewed daily for 5 days from day 1 to day 5. The control group of no-drug was set up. ZF were cultured in 35°C. Survival analysis was done, and MTCs were calculated. The MTC of each drug was defined as the highest concentration without ZF death.

### 4. Drug efficacy assessment in *M. abscessus*-infected ZF

3dpf ZF with homogeneous distribution of *M. abscessus* were selected, and randomly placed into 24-well plates, with each well of having 1ml water and 10 ZF. In preliminary experiments, noninfected embryos were exposed to increasing concentrations of MFX and AZM, and observed under a microscope. The drug concentration with no signs of toxicity-induced killing or developmental abnormalities were tested. Different doses were tested, corresponding to 31.25×, 62.5×, 125×, 250× and 500× the MIC of MFX, and 3.9×, 7.8×, 15.625×, 31.25× and 62.5× the MIC of AZM, based on the values determined using the microdilution method. Finally, with MFX concentrations of 62.5μg/ml, 125μg/ml, 250μg/ml, 500μg/ml, 1000μg/ml, and AZM of 15.625μg/ml, 31.25μg/ml, 62.5μg/ml, 125μg/ml, 250μg/ml, separately in each well, no ZF death appeared. The maximum concentrations tested in the next process should below the MTC. 20 3dpf ZF in each concentration were tested. Drug-containing water was renewed daily for 5 days from day 1 to day 5 post infection. The control group without drug were set up for control. ZF were cultured in 35°C. Survival curves were determined by recording dead ZF every day.

3 days after infection, 5 ZF in each concentration were collected and pictured. Fluorescence microscopy of infected ZF was performed using Nikon NIS-Elements D 3.10 fluorescence microscope. Final image analysis and visualization were performed using GIMP 2.6 freeware to merge fluorescent and differential inference contrast (DIC) images and to adjust levels and brightness and to remove out-of-focus background fluorescence. Images of fluorescence intensity in different concentration were calculated by pixel.

3 days after infection, 5 ZF in each concentration were pictured, and the inhibition rate in each concentration was calculated. Inhibition rate (%) was calculated by formula: inhibition rate (%) = (S_control group_-S_drug group_)/S_control group_×100% (S: fluorescence intensity by pixel).

*M. abscessus* may disseminated in hear, brain, vein, liver and eyes. In order to analyze drug efficacy against *M. abscessus* dissemination, fluorescence dissemination under different drug concentration in ZF were observed, pictured and analyzed.

From day 1 to day 3, 5 ZF in each concentration were collected, lysed individually in 2% Triton X-100-PBS, and resuspended in PBS with Tween 80. Several 10-fold dilutions of homogenates were plated on 7H10 containing 500 mg/liter hygromycin and BBL MGIT PANTA (Becton Dickinson), used as recommended by the supplier. Colony-forming unit (CFU) were enumerated after 4 days of incubation at 35°C. Results are expressed as mean log10 CFU per ZF.

### 5. Statistical analyses

Statistical analyses of comparisons between Kaplan-Meier survival curves were performed using the log-rank test with SPSS software. CFU counts and quantification experiments were analyzed using one-way analysis of variance and Fisher’s exact test, respectively. Statistical significance was assumed at P values of 0.05.

## Result

### 1. MICs

After DST *in vitro*, the MFX MIC against *M. abscessus* standard isolate showed 2μg/ml, meaning moderately susceptible to MFX.

### 2. Microinjection of *M. abscessus* into ZF

Different bacteria concentrations and observation duration were tested for establishing infected ZF model. High concentration would lead to rapid death. By contrary, low concentration would not achieve enough fluorescence imaging pictured by microscope. Three *M. abscessus* concentration of 1.6×10^9^/ml, 2×10^9^/ml and 5×10^9^/ml were chosen for testing. The bacteria amount of 12800, 6400, 3200, 1600 and 800 were tested for injection. Observation duration from 3 to 7 days were tested. Finally, *M. abscessus* concentration of 5×10^9^/ml with the amount of 1600 *M. abscessus* were injected into ZF, and the observation duration of 5 days was chosen.

### 3. MTC of MFX and AZM in ZF model *in vivo*

Each drug’s MTC was tested (table 1). The concentrations that couldn’t influence ZF survivability could be selected for the next process. MFX with concentration of ≤1000μg/ml and AZM of ≤ 250μg/ml would not impact ZF survivability. Finally, the MFX concentration of 62.5μg/ml, 125μg/ml, 250μg/ml, 500μg/ml, 1000μg/ml, and AZM concentration of 15.625μg/ml, 31.25μg/ml, 62.5μg/ml, 125μg/ml, 250μg/ml were chosen for the next process.

**Table 1:**
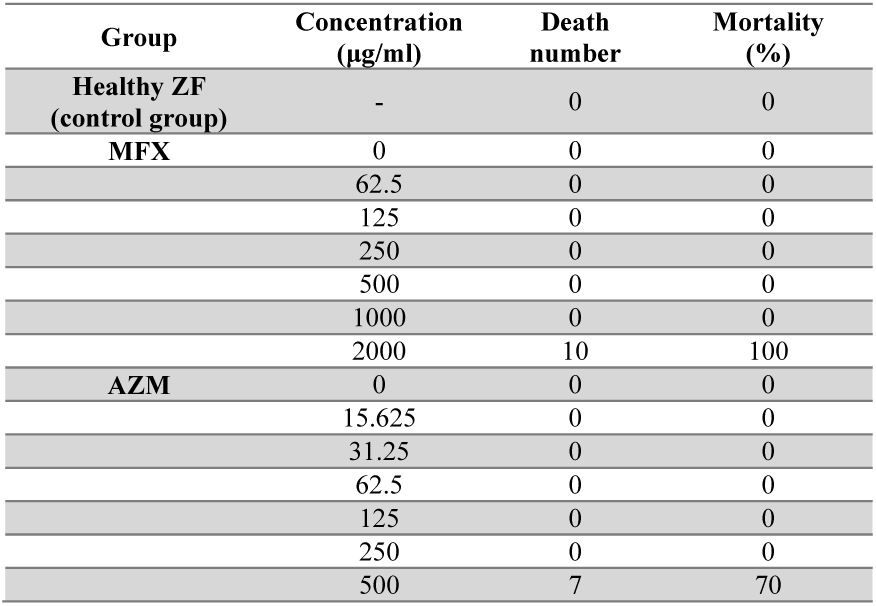
ZF survivability in different concentrations of MFX and AZM (n=10 of each group)

| Group | Concentration (µg/ml) | Death number | Mortality (%) |
| --- | --- | --- | --- |
| **Healthy ZF (control group)** | - | 0 | 0 |
| **MFX** | 0 | 0 | 0 |
|  | 62.5 | 0 | 0 |
|  | 125 | 0 | 0 |
|  | 250 | 0 | 0 |
|  | 500 | 0 | 0 |
|  | 1000 | 0 | 0 |
|  | 2000 | 10 | 100 |
| **AZM** | 0 | 0 | 0 |
|  | 15.625 | 0 | 0 |
|  | 31.25 | 0 | 0 |
|  | 62.5 | 0 | 0 |
|  | 125 | 0 | 0 |
|  | 250 | 0 | 0 |
|  | 500 | 7 | 70 |

### 4. Drug efficacy assessment in *M. abscessus*-infected ZF

#### 4.1 ZF survival

We tested a wide range of AZM concentrations from 15.625μg/ml to 250μg/ml, and MFX concentration ranging from 62.5μg/ml to 1000μg/ml. Supplementing the ZF-containing water with these drug concentrations led to no toxicity, as measured by pre-experiment.

When infected ZF were exposed for more than 2 days to the concentrations of AZM tested, a significant increased survival rate (P=0.000) was observed between different AZM concentration (Figure 1A). And increased survival was associated with high dose of AZM. Exposure to a higher dose of AZM further extended the life span of infected ZF. The treatment with the low AZM doses failed to restrict mycobacterial growth. This result shows that AZM had significant activity against *M. abscessus in vivo* and is efficient in this ZF test system against *M. abscessus* infection. However, although some restriction to mycobacterial growth by MFX was observed, the association between increased survival and high dose of MFX is not significant (figure 1B). With the MFX concentration increased, the survival curve didn’t show significant decrease between different MFX concentration (P=0.267).

**Figure 1A:**
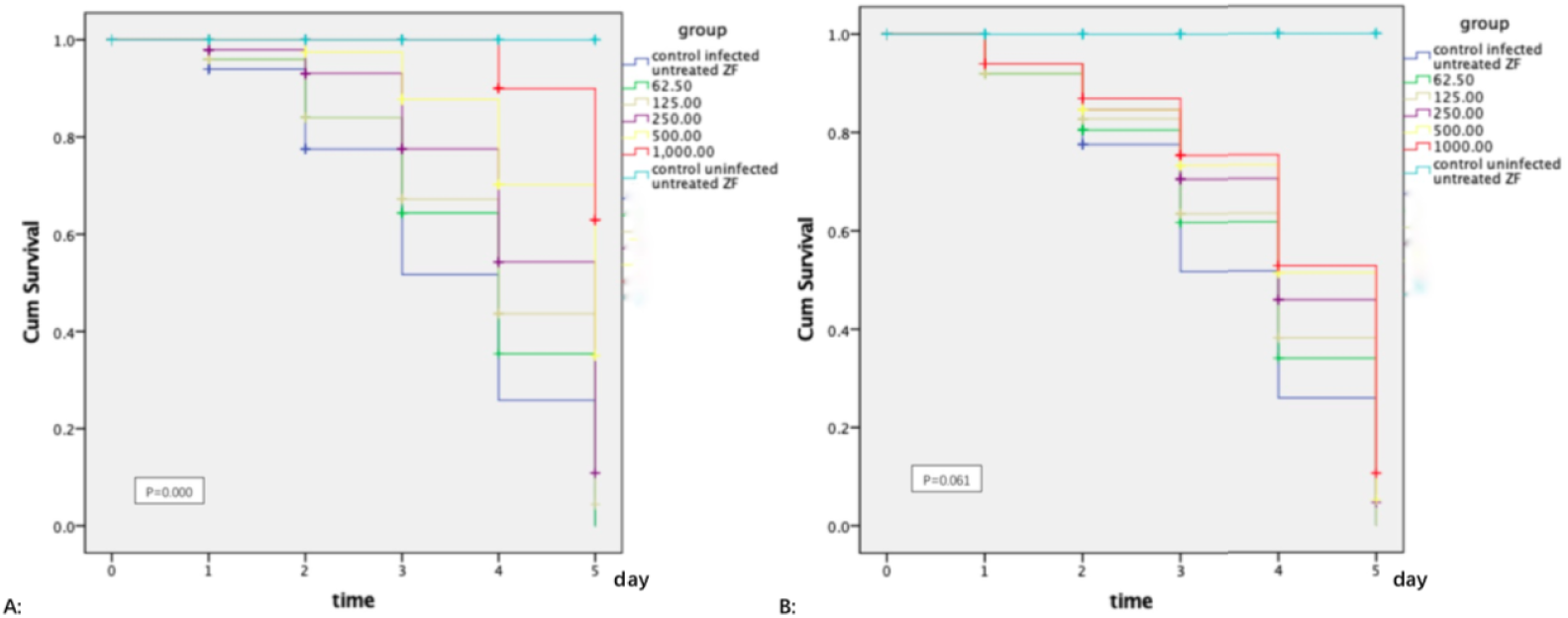
Increased survival was associated with high dose of AZM. The treatment with the low AZM doses failed to restrict mycobacterial growth. The survival curve showed significant difference between difference AZM concentration group (P=0.000). Figure 1B: Although some restriction to mycobacterial growth by MFX was observed, the association between increased survival and high dose of MFX is not significant (P=0.267). The statistical comparison was tested between different drug concentration but without control uninfected untreated ZF.

#### 4.2 Bacterial burdens

Drug efficacy against bacterial burden by CFU loads was also analyzed. Increased AZM concentration was associated with lower bacterial burdens as determined quantitatively by CFU plating (figure 2A). The treatment with lower doses restricted less mycobacterial growth. The same trend was observed in MFX. The MFX concentration correlated with CFU loads (figure 2B). In both AZM and MFX groups, no significant difference was observed between different concentration group (P > 0.05).

**Figure 2A:**
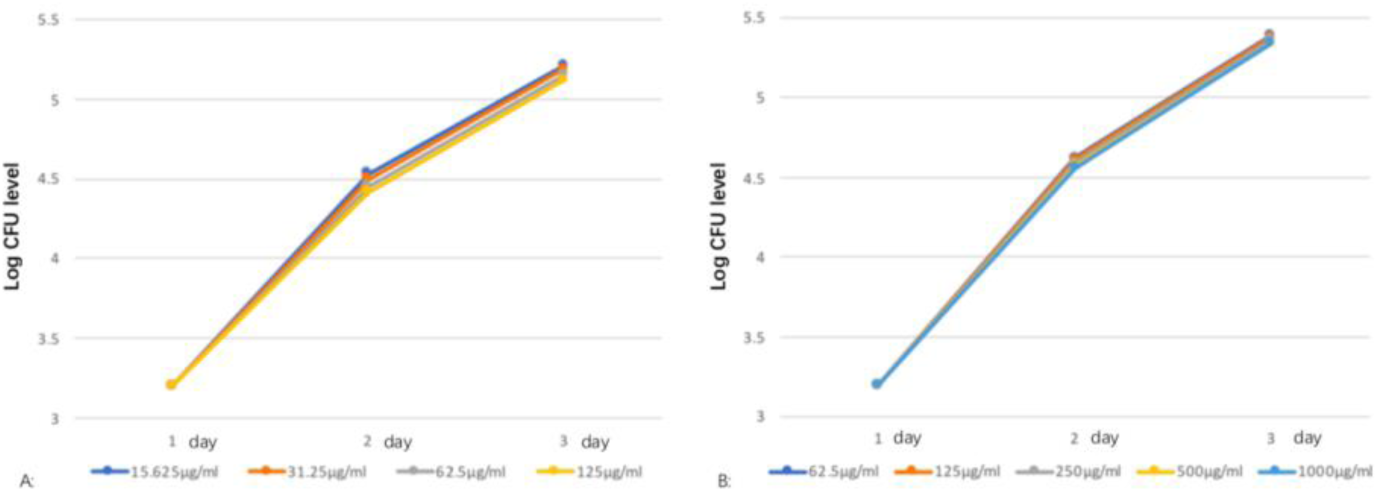
From day 1 to day 3, 5 ZF in each concentration were collected, lysed and plated on 7H10. Increased AZM concentration was associated with lower bacterial burdens as determined quantitatively by CFU plating. The treatment with lower doses restricted less mycobacterial growth. No significant difference was observed between different AZM concentration group (P > 0.05). Figure 2B: The same trend was observed in MFX. The MFX concentration correlated with CFU loads. No significant difference was observed between different MFX concentration group (P > 0.05).

#### 4.3 Bacterial fluorescence intensity in ZF

3 days after infection, 5 ZF in each concentration were collected and pictured. In our study, exposure to AZM was associated with a significant reduction in the numbers of abscesses (figure 3A). Under different AZM concentration (15.625μg/ml, 31.25μg/ml, 62.5μg/ml, 125μg/ml), bacterial fluorescence intensity in ZF showed significant decrease (161828 ± 6605, 157329 ± 5356, 142300 ± 13715, 132942 ± 11243) (figure 3A). This decrease of fluorescence intensity was consistent with the inhibition rate (figure 4A). The inhibition rate also showed significant difference when comparing with no drug group (P<0.05), which means AZM showed good inhibition efficacy. However, exposure of infected ZF to MFX showed no significant decrease in the frequency of abscesses (figure 3B). With MFX concentrations increased, fluorescence intensity decreased slightly under fluorescence microscope. In MFX concentrations of 62.5μg/ml, 125μg/ml, 250μg/ml, 500μg/ml and 1000μg/ml, fluorescence intensities in ZF were 247306, 243523, 229586, 221573 and 219640 pixels (figure 3B), and the inhibition rates were 0%, 1%, 7%, 10% and 11% respectively, with all P value >0.05 when comparing with control group (figure 4B).

**Figure 3A:**
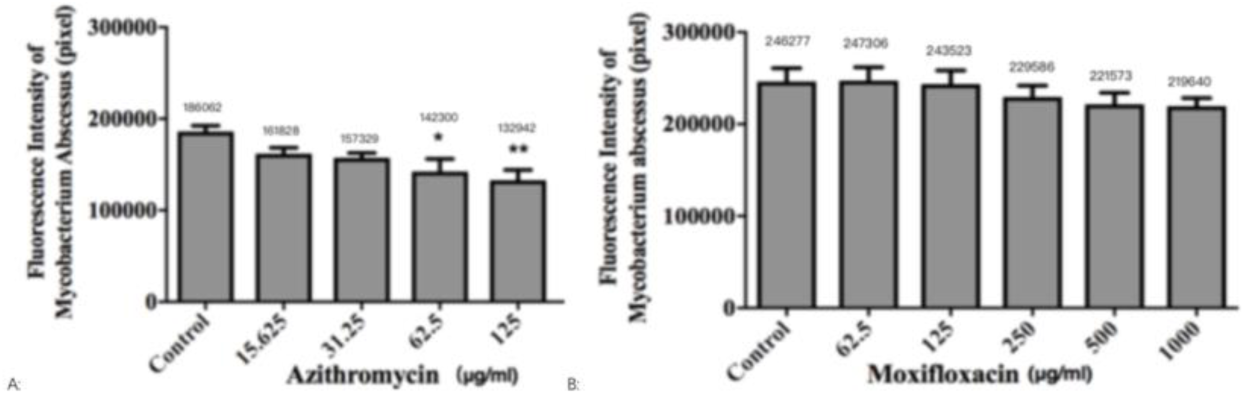
3 days after infection, 5 ZF in each concentration were collected and pictured. The fluorescence intensity (by pixel) of *M. abscessus* in different concentrations of AZM (figure 4B) was compared with control group without drug. Figure 4B: The fluorescence intensity of *M. abscessus* in different concentrations of MFX was compared with control group without drug. N=20 for each group. . means P <0.05.

**Figure 4A:**
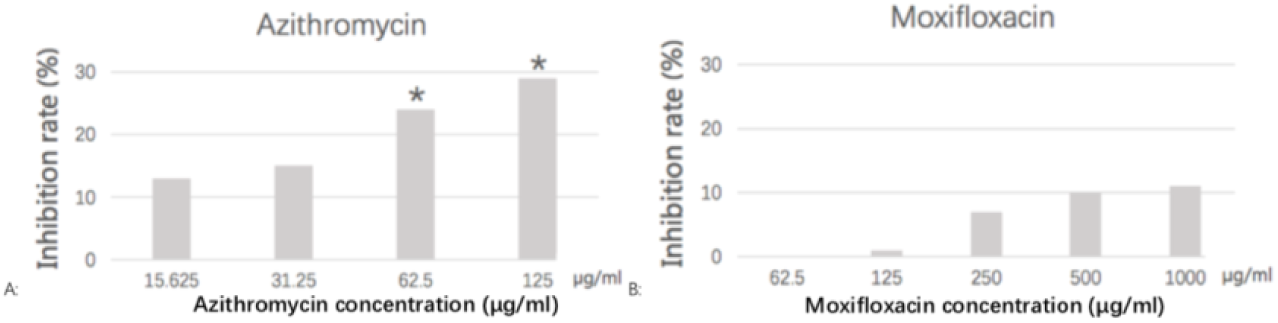
3 days after infection, 5 ZF in each concentration were pictured, and the inhibition rate in each concentration was calculated. The AZM inhibition rates in concentration 15.625μg/ml, 31.25μg/ml, 62.5μg/ml and 125μg/ml were 0%, 13%, 15%, 24% and 29% respectively, with all P value >0.05 when comparing with control group. Figure 4B: The MFX inhibition rates in concentration 62.5μg/ml, 125μg/ml, 250μg/ml, 500μg/ml and 1000μg/ml were 0%, 1%, 7%, 10% and 11% respectively, with all P value >0.05 when comparing with control group. One-way analysis of variance and t test was calculated. . means P <0.05.

#### 4.4 Drug effect on bacteria dissemination

AZM and MFX’s effects on bacteria fluorescence dissemination were analyzed. In AZM control group without drug, *M. abscessus* disseminated in heart, brain and vein. The transferring occurrence rate was 50%. In AZM 15.625μg/ml group, *M. abscessus* disseminated in brain and vein, with transferring occurrence rate of 30%. In AZM 31.25μg/ml, 62.5μg/ml and 125μg/ml group, *M. abscessus* disseminated in only vein, with transferring occurrence rate of 20%. All the transferring rates in different concentrations were compared with control group with p > 0.05. In MFX control group without drug, *M. abscessus* disseminated in liver, heart, brain and vein. The transferring occurrence rate was 70%. In MFX 62.5μg/ml group, *M. abscessus* disseminated in heart and vein, with transferring occurrence rate of 60%. In MFX 125μg/ml group, *M. abscessus* disseminated in brain and vein, with transferring occurrence rate of 50%. In MFX 250μg/ml, 500μg/ml and 1000μg/ml group, *M. abscessus* disseminated in brain and vein, with transferring occurrence rate of 40%. All the transferring rates in different concentrations were compared with control group with p > 0.05. Although both two groups showed some inhibition on *M. abscessus* dissemination, no significant difference was observed in both AZM and MFX groups when compared with control group.

Together, all these results above suggest that AZM exerts a therapeutic effect by preventing the development of abscesses and protecting the ZF from bacterial killing. However, MFX exerts limited effect by preventing the development of abscesses and protecting the ZF from bacterial killing.

## Discussion

Animal models are currently limited to study pathogenesis unless very large doses of bacilli are given intravenously. If small doses are given, there is little evidence that a productive infection is even fully established. Consequently, better models are required to elucidate pathogenesis and to enable new drugs for M. abscessus infections to be tested. This led to the recent development of the zebrafish model to assess the suitability and sensitivity of clinically relevant drugs in M. abscessus-infected embryos [19,25,27,28]. Small bacterial doses can be used in this model to allow visualizing, in a dose- and time-dependent manner, the dynamics of infection and physiopathological markers, such as cords and abscesses, in the presence of an active compound [19]. The injection of a small inoculum allows administration of homogenous bacterial suspensions without obstructing the needle during the microinjection procedure. In this study, we evaluated the *in vivo* drug activity by ZF model.

Recognized as a cause of chronic pulmonary infections, especially in individuals with altered host defenses or disrupted airway clearance mechanisms, *M. abscessus* appears as a major infectious threat to the airway in CF patients, and reports suggest increased prevalence in recent years [19,29]. This situation is worsened by the fact that antibio-therapy against *M. abscessus* is often unsuccessful and/or poorly tolerated by patients. *M. abscessus* is notorious for being intrinsically resistant to most antibiotics [7], thus rendering these infections particularly complicated, difficult to treat, and associated with a high rate of therapeutic failure [30].

Some studies showed that MFX had less or none activity against *M. abscessus in vitro* [15-18]. However, some other studies showed that MFX had good activity *in vitro* against *M. abscessus* and was recommended as one of the antibiotic regimens for adults with *M. abscessus* disease [12, 14]. There is not enough recommendation *in vivo* for the preferential use of this drug. As a result, work should address the *in vivo* efficacy.

We have previously shown that MFX with a moderate but significant inhibition efficacy on M. abscessus *in vitro* [14]. In this study, we evaluated the *in vivo* activity of MFX against *M. abscessus. M. abscessus* exists as two variants: rough and smooth. Ex vivo and *in vivo* studies have described the hypervirulence phenotype of the R versus the S morphotype [31,32], and epidemiological studies have confirmed the persistence and acute respiratory syndromes caused by the R morphotype [33-35]. The major difference between the R and S variants is the loss of a surface-associated glycopeptidolipid (GPL) [36]. In this study, we chose S morphotype for testing, which covered 53% of the whole *M. abscessus* in China [37].

In agreement with a recent study addressing the activity of MFX against several NTMs [14], we found that MFX exhibited low MIC values against standard *M. abscessu* isolate *in vitro*. The efficacy of MFX against *M. abscessus* was also investigated by monitoring the survival and bacterial burden of infected ZF treated with MFX. AZM had an excellent activity on *M. abscessus* and was tested together with MFX for comparation. Fluorescenece intensity of *M. abscessus* amount under different concentrations in ZF was analyzed. AZM showed good activity for decreasing bacteria amount in ZF, which further verified our experiment’s stability and accuracy. However, MFX showed no significant decrease in bacteria inhibition compared with control group. With the MFX concentration increasing, although we could see mild decreasing of bacteria fluorescence intensity, there still showed no significant difference compared with control group. MFX’s inhibition rate on *M. abscessus* was calculated. AZM showed significant decrease on inhibition rate when compared with control group. While no significant difference on inhibition rate was observed in MFX group when compared with control group. MFX’s effect on ZF survival was also analyzed. In the ZF model, AZM increased the survival of the ZF as previously reported. And the statistical analysis did demonstrate a significant difference compared with no drugs. However, the efficacy of MFX is likely to be poor since the Kaplan-Meier survival curve did not show significant inhibition on infected ZF. These results above all verified that MFX may have limited activity against *M. abscessus in vivo* especially compared with AZM.

By analyzing bacterial dissemination and CFU loads, it showed no significant result even in AZM group. Although some inhibition of MFX and AZM on *M. abscessus* dissemination in ZF was observed, there was no significant difference when compared with control group. Same result was also observed in analyzing the drug efficacy on CFU loads. Although positive impact on inhibiting CFU was observed, there was no significant result even. Although the CFU was recommended for ZF model [19,24], in our study it is useless for assessing drug activity *in vivo* using ZF model. Experimental method may need to be improved.

To sum up, we report here a robust and sustained effect of MFX in infected zebrafish. Our results suspect the efficacy of MFX *in vivo* in this animal model, allowing visualizing in a dose- and time-dependent manner the dynamics of cord and bacterial fluorescence and loads. MFX exerted a limited impact on ZF survival. This comports well with failure of the MFX-containing regimens in clinical practice [38]. But such confirmation will require multivariable pharmacometric analyses of clinical data, which do not currently exist for MFX and *M. abscessus* pulmonary disease [39]. Indeed, no such data exist, as far as we know, for any of the drugs used in treatment of pulmonary *M. abscessus*, since there is a lack of clinical trials and large prospective clinical cohort studies for this disease [38].

Further more, the present study reports the usefulness of ZF as a preclinical model to evaluate in real time the efficacy of MFX and AZM against *M. abscessus* infection. Since ZF have been successfully used in the past to test the efficacy of three clinically relevant drugs, clarithromycin, imipenem and bedaquiline [19,24,25], future studies should address the *in vivo* efficacies of other drugs using the zebrafish model of *M. abscessus* infection.

There are limitations remained: 1. Small sample size limited the appearance of drug efficacy. 2. We only test standard isolate ATCC19977. More works should be further exploited to compare the intrinsic activity of antibiotics *in vivo* in ZF infected with the three subspecies of the *M. abscessus* complex, *M. abscessus* sensu stricto, *Mycobacterium massiliense*, and *Mycobacterium bolletii*, which are known to respond differently to antibiotics *in vitro*. 3. Due to these strain-to-strain variations, clinical strains should also be tested and may greatly help the clinician to select optimal drug treatments. 4. The time-course of death induced by *M. abscessus* is so rapid, with up to 50% of ZF dying 5 dpi and 100% within 10 dpi. So enough observation on ZF survival is limited.

## Acknowledgments

This work was supported by Beijing Hospital Authority Youth Programme [QML20171602], Tongzhou District’s Two Supreme Talent [YH201911], Key laboratory of capital medical university open research project and Beijing Tuberculosis & Thoracic Tumor Research Institute cultivation project & 13th Five National Major Scientific and Technological Projects [2017ZX09304009].

